# Plus-end directionality is present in the catalytic core of kinesin-14 minus-end directed motors

**DOI:** 10.1101/292375

**Authors:** M. Yamagishi, J. Yajima

## Abstract

Molecular motors move along microtubules unidirectionally. However, the origin of directionality is not clear. Here kinesin-14 monomers were engineered such that they could be anchored via either their N- or C-termini, leaving the opposite terminal regions mechanically disconnected from the surface. We find that the conserved catalytic motor core of kinesin-14 has intrinsic plus-end directionality and that the proper function of the neck-helix region is required to achieve minus-end directionality.

---

The uni-directional movement of kinesin motor proteins along microtubules is important in many cellular processes in eukaryotes, including organelle transport and cell division. N-kinesins (kinesin-1 to 12) with a motor domain at the N-terminus move towards the plus-ends of microtubules, whilst C-kinesins (kinesin-14) with a motor domain at the C-terminus move towards the minus-end. Although N-kinesin and C-kinesin have a similar catalytic core structure, these kinesins have their own unique structures at either their N- or C-terminus (**Fig. 1*A***, **Fig. S1*A*** and ***B***). N-kinesin has a C-terminal neck including a 15 amino acid region termed the neck-linker (1, 2). C-kinesin has an N-terminal coiled-coil helix termed the neck-helix (3) and a C-terminal 15 amino acid region termed the neck-mimic (4).

**Figure 1.**
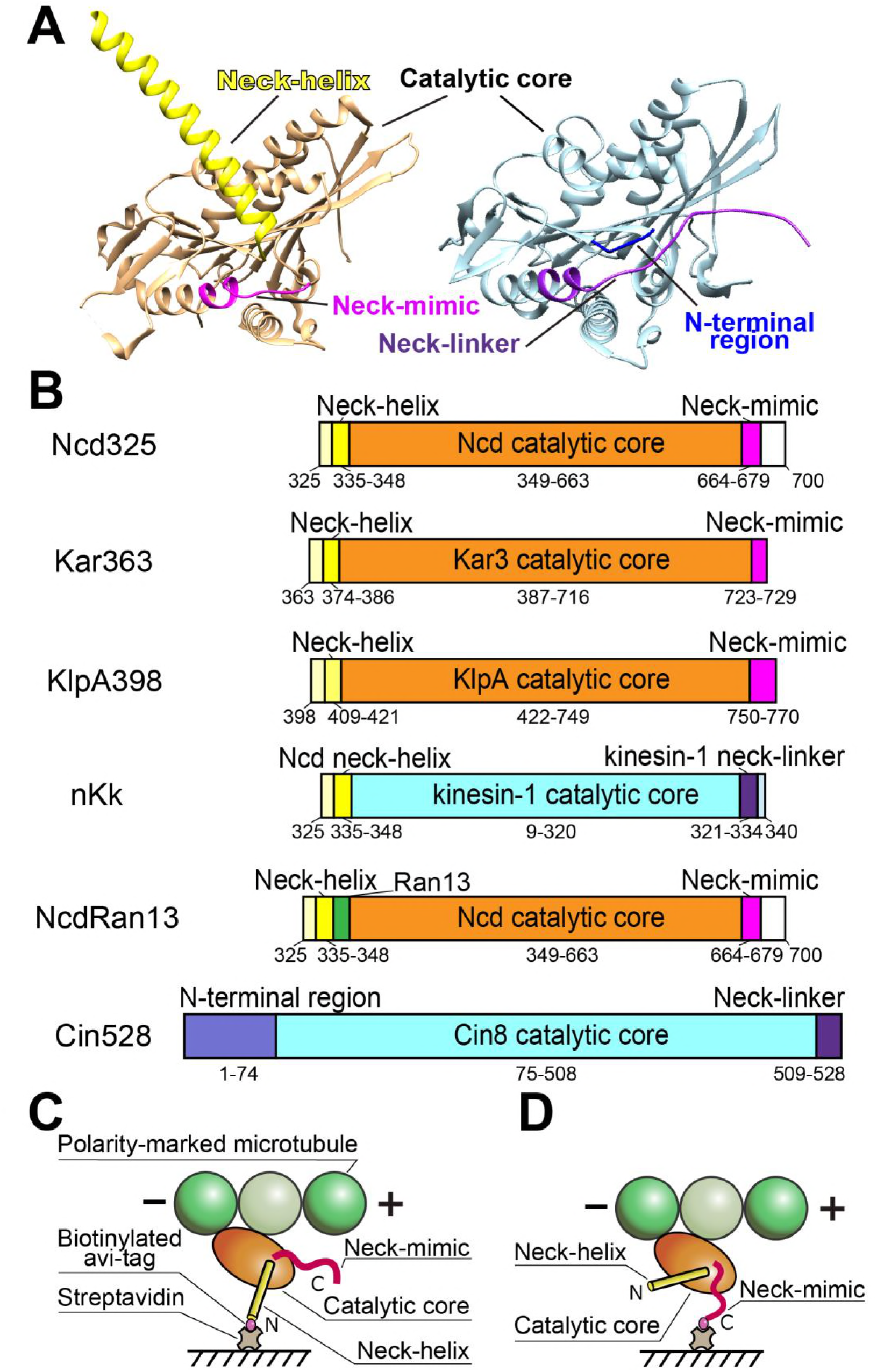
Observation of the directionality of N- or C-linked monomeric kinesin-14. (*A*) The 3D structure of Ncd (PDB: 3U06, chain B) and kinesin-1 (PDB: 4HNA). The catalytic core (Ncd, orange; kinesin-1, cyan) has high structural homology among kinesins. The N-terminal neck-helix of Ncd (yellow) and the N-terminal region of kinesin-1 (blue) have little structural homology. The C-terminal neck-mimic of Ncd (magenta) and the neck-linker of kinesin-1 (purple) have slight structural homology. (*B*) N- or C-linked kinesin monomeric constructs. All constructs (kinesin-14s, kinesin-1– Ncd chimera, Ncd mutant and kinesin-5 Cin8) have biotinylated peptide (avi-tag) at either their N-terminus or C-terminus. Apart from nKk, all constructs use the native neck-helix, N-terminal region, neck-mimic or neck-liner for the corresponding kinesin motor. NcdRan13 constructs have an insertion of random 13 residues ESGAKQGEKGESG (green) between neck-helix and catalytic core. (*C* and *D*) Scheme of a single headed kinesin-14 anchored to the streptavidin-coated substrate via its N-terminus (*C*) and C-terminus (*D*), respectively.

In an effort to identify determinants for the direction of movement of kinesins along microtubules, several groups have constructed chimeras. A chimera consisting of a C-kinesin (Ncd) (5) neck-helix and neck-mimic regions were fused to an N-kinesin (kinesin-1) catalytic motor core region at the N- and C-termini, respectively. The chimeras, which were designed to form a dimer (NcdKHC1) (6) or monomer (nKn664) (7), have minus-end directionality, reversing the polarity of kinesin-1 movement (**Fig. S1*C* and *D***). A further finding was that chimeras with mutations in either the N-terminal neck-helix – catalytic core junction (NcdKHC5) (6) or catalytic core – C-terminal neck-mimic junction (nKn669) (7) retain the plus-end directionality of the kinesin-1 (**Fig. S1*E*** and ***F***). These results indicate that proper functioning of both Ncd neck-helix and neck-mimic are required to reverse the plus-end polarity of kinesin-1 movement. These chimeras also had neither a kinesin-1 neck-cover strand (8) nor kinesin-1 neck-linker, further suggesting that the plus-end polarity determinants are present in the kinesin-1 catalytic core itself.

Other studies have shown that complementary chimeras consisting of the Ncd catalytic core region fused to a kinesin-1 neck-linker and coiled-coil region, ncd-Nkin (9) and NK-1 (2), moved towards the microtubule plus-end, reversing the minus-end polarity of Ncd movement (**Fig. S1*G*** and ***H***). Another chimera, consisting of the Ncd catalytic motor core region fused to a kinesin-1 neck-linker without the dimerizing region at the C-termini and fused to the Ncd neck-helix with the dimerizing region at the N-termini, did not invert the minus-end polarity of Ncd movement (NcdKHC6) (6) (**Fig. S1*I***). These results were interpreted as indicating that a positional bias of the microtubule-unattached motor domain of two-headed kinesin, where the unattached-motor domain has been reported to be tilted towards the microtubule plus- or minus-end for plus- or minus-end directed kinesins respectively (10), determined the directionality. Sablin *et al.* also demonstrated that a two-headed Ncd mutant, in which the neck-helix was mutated by randomizing 12 residues, showed plus-end-directed motility (ncd-ran12) (11) (**Fig. S1*J***). This study suggested that either (i) dysfunction of the neck-helix caused the unattached motor domain of two-headed Ncd to tilt towards the microtubule plus-end or (ii) the Ncd motor domain with dysfunction of the neck-helix contains a plus-end determinant, like the kinesin-1 catalytic core (12).

To distinguish between these possibilities, we tested if the catalytic core of C-kinesins possesses a plus-end directionality and if this directionality is affected by the geometry of attachment of the motor head. We made minimal motor domain constructs comprising different kinesin-14 single heads (*Drosophila* Ncd (5); 325–700 aa, *Saccharomyces cerevisiae* Kar3 (13); 363–729 aa and *Aspergillus nidulans* KlpA (14); 398–770 aa) fused to a biotinylated peptide (avi-tag) (15) at either their N-terminus (BP-Ncd325, BP-Kar363 and BP-KlpA398) or C-terminus (Ncd325-BP, Kar363-BP and KlpA398-BP) (**Fig. 1*B***). These constructs allowed us to bind kinesin-14 monomers to a streptavidin-coated substrate via either their N-termini (neck-helix) or C-termini (neck-mimic) (**Fig. 1*C*** and ***D***). For the N-linked kinesin-14 monomers (BP-Ncd325, BP-Kar363 and BP-KlpA398) attached to the streptavidin-coated substrate via the biotinylated avi-tag fused to their N-termini, the N-terminal neck-helix is connected to the substrate so that it can transmit the microtubule sliding force (**Fig. 1*C***). For the C-linked kinesin-14 (Ncd325-BP, Kar363-BP and KlpA398-BP) attached to the substrate via their C-terminal neck-mimic, the distal end of the N-terminal neck-helix is disconnected and is hanging free in solution (**Fig. 1*D***).

We first assayed the directionality using an *in vitro* polarity-marked microtubule sliding assay (**Fig. 2*A***) (7, 16). N-linked kinesin-14s showed minus-end directionality (**Fig. 2*B*, *D*** and ***F***, Table 1, see also **Movie 1 in the Supporting Material**), which is consistent with previous reports (3, 13, 14). Remarkably, we found that surface of C-linked kinesin-14 drove microtubule sliding with the bright minus-ends leading, indicating a plus-end-directed motor activity, like kinesin-1 (**Fig. 2*C*, *E*** and ***G***, **Table 1**, see also **Movie 1 in the Supporting Material**). Sliding velocity driven by each of the three C-linked kinesin-14 monomers is 2~10 times faster than that of the corresponding N-linked kinesin-14 monomers (3) (**Table 1**). These data clearly indicate that directionality in kinesin-14 monomer driven microtubule sliding assays depends on which kinesin-14 terminus is tethered to the surface. Activity of kinesin-14’s catalytic motor core coupled via the C-terminal neck-mimic produces plus-end directionality whilst coupling via the N-terminal neck-helix produces minus-end directionality. This extends an earlier prediction that the function of the C-terminal neck-mimic is similar to that of the kinesin-1 neck-linker (4, 12, 17, 18) by showing that the neck-mimic can effectively transmit the plus-end biased conformational changes that occur in the kinesin-14’s catalytic core when the neck-mimic is used to couple the catalytic motor core to the glass surface.

**Table 1.**
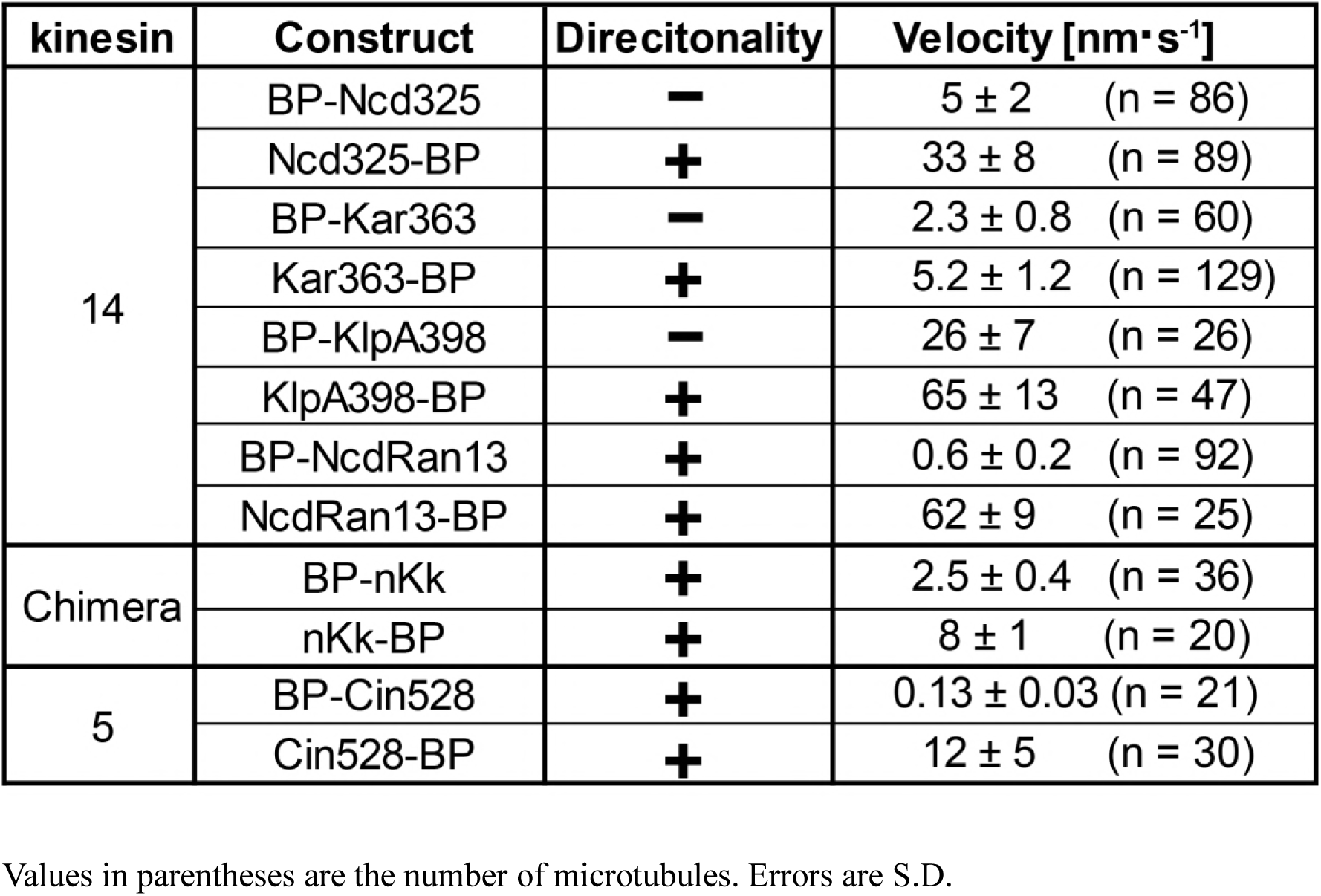
The direction and velocity of monomeric kinesins

**Figure 2.**
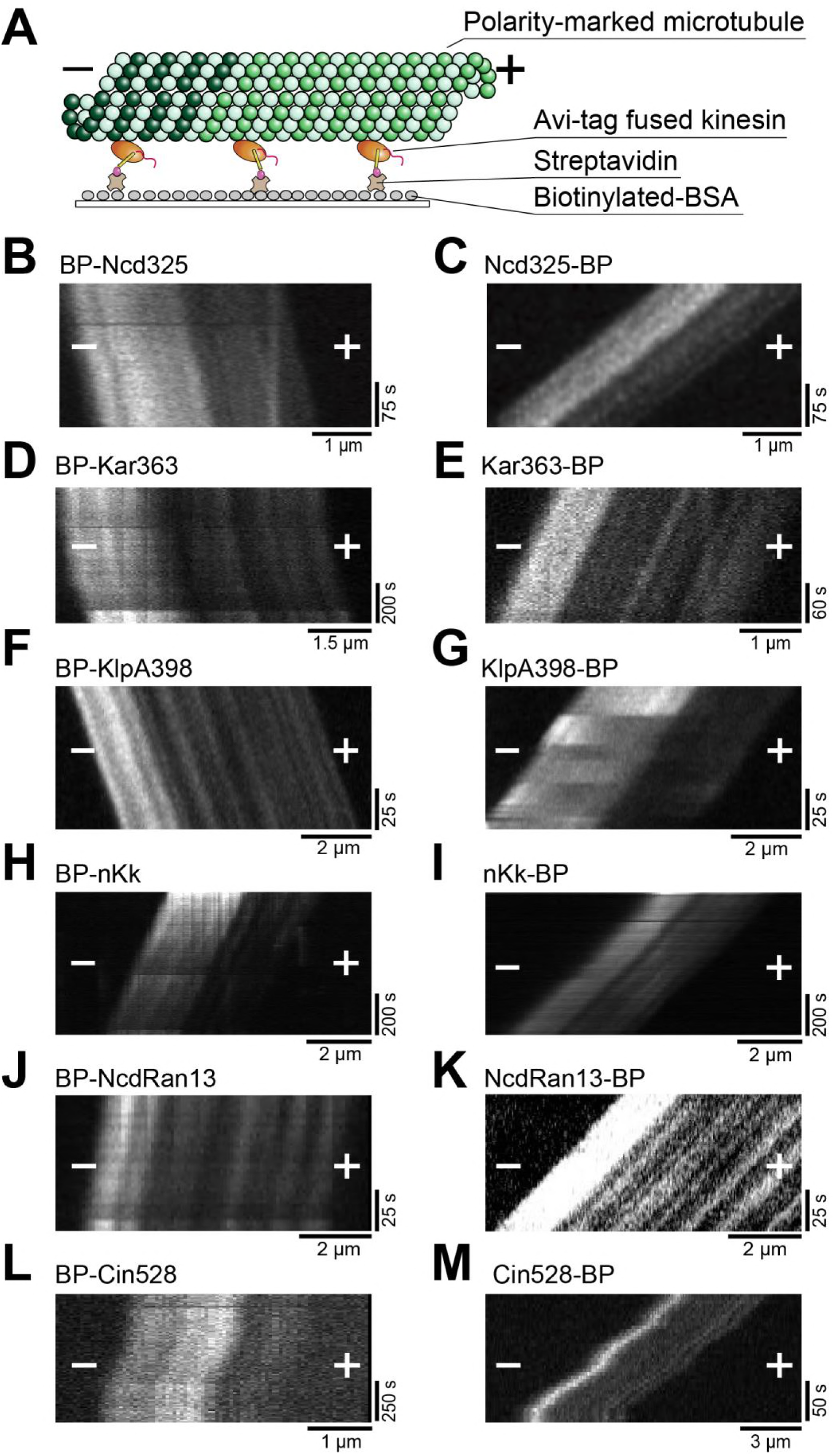
Polarity-marked microtubule sliding assays driven by N- or C-linked monomeric kinesins. (*A*) Scheme of the *in vitro* polarity-marked microtubule sliding assay. Single-headed kinesins fused to a biotinylated tag (avi-tag) are anchored to biotinylated BSA via biotin-streptavidin linkage. (*B*~*G*) Typical kymographs of the polarity-marked microtubule sliding driven by N-linked monomeric Ncd (*B*), Kar3 (*D*), KlpA (*F*) and C-linked monomeric Ncd (*C*), Kar3 (*E*), KlpA (*G*) are shown, respectively. N-linked monomeric kinesin-14s slide microtubules with their plus-ends leading, indicating minus-end-directed motor activity, whereas C-linked monomeric kinesin-14s slide with their minus-ends leading, indicating plus-end-directed motor activity. (*H*~*M*) Typical kymographs of the polarity-marked microtubules sliding driven by N- or C-linked monomeric chimera kinesin nKk (*H* and *I*), Ncd mutant NcdRan13 (*J* and *K*) and the kinesin-5 Cin8 (*L* and *M*) are shown. Both N- and C-linked monomeric kinesins slide microtubules with their minus-ends leading, indicating plus-end-directed motor activity. The plus (+) and minus (-) signs refer to the plus-end and minus-end of the microtubules, respectively.

The N-terminal neck-helix outside of the catalytic motor core appears to have a key role in generating minus-end directionality. However, previous studies reported that Ncd – kinesin-1 chimeras, which included both the Ncd neck-helix and neck-mimic regions, showed minus-end directionality, reversing the plus-end directionality of the kinesin-1 motor (NcdKHC1 (6) and nKn664 (7), see also **Fig. S1**). To determine if the Ncd neck-helix alone can reverse kinesin-1 directionality in the absence of the Ncd neck-mimic, we made a monomeric Ncd – kinesin-1 chimera, nKk, composed of the Ncd neck-helix followed by the kinesin-1 catalytic motor core and kinesin-1 neck-linker in both the N- and C-linked configurations (**Fig. 1*B***). We found that both N- and C-linked nKk showed plus-end motility (**Fig. 2*H*** and ***I***, **Table 1**, see also **Movie 1 in the Supporting Material**), showing that the Ncd neck-helix alone is not sufficient to reverse the directionality of the kinesin-1 catalytic motor core and that reversing the directionality of the kinesin-1 motor core require the neck-helix – neck-mimic interaction. To further investigate the role of the N-terminal neck-helix in determining the minus-end directionality, we made a monomeric Ncd mutant, NcdRan13, in which 13 randomized residues were inserted between the neck-helix and catalytic core, in both the N- and C-linked configurations (**Fig. 1*B***). NcdRan13 is the monomeric version of dimeric ncd-ran12 reported as a plus-end-directed motor (11) (**Fig. S1*J***) and differs from ncd-ran12 by insertion of 13 amino acids in the junction. Microtubule sliding assays of both N- and C-linked NcdRan13 showed plus-end movement of the motor on microtubules with a velocity of 0.6 ± 0.2 nm s^−1^ (mean ± SD, n = 92) and 62 ± 9 nm s^−1^ (mean ± SD, n = 25), respectively (**Fig. 2*J*** and ***K***, **Table 1**, see also **Movie 1 in the Supporting Material**). This finding, together with the plus-end directionality of nKk in the absence of the neck-mimic region, shows that coupling of the neck-helix to the motor core or the neck-mimic is critical to generate minus-end directionality and that in the absence of this precise coupling the motor generates plus-end directionality.

Recently, the kinesin-14 KlpA has been shown to switch its directionality depending on whether KlpA works as single molecule or in a team (14). Motor number dependent directionality was also reported for Cin8 from the kinesin-5 *Saccharomyces cerevisiae* (19, 20). However, it was not tested if the directional switching also occurred with the monomeric constructs. To test this, we examined the dependence of directionality on the motor number of monomeric KlpA in both the N- and C-linked configurations. We found that the directionality of both N- and C-linked KlpA did not change in a polarity-marked microtubule sliding assay as the monomeric concentrations ranged from 0.25 to 1 μM, whereas the sliding velocity of N-linked KlpA slightly changed (**Fig. 2*F* and *G***, **Fig. 3** *red*). We also found that both N- and C-linked monomeric Cin8 did not change directionality with changes in motor number (**Fig. 2*L*** and ***M***, **Fig. 3**, *blue*). This indicates that the motor number of monomeric constructs is not relevant in determining directionality, which is consistent with a recent report on Cut7, the kinesin-5 from *Schizosaccharomyces pombe* (21). Therefore, the directionality is determined by the end used for anchoring the monomeric constructs. We also assayed the motor directionality of kinesin-14 monomeric constructs (KlpA398 and Ncd325) by attaching them to quantum dots (QDs) and allowing them to move along immobilized microtubules (**Fig. 4*A***). QDs carrying either N- or C-linked constructs did not change directionality (**Fig. 4*B***~***E***, **Table 1**, see also **Movie 2 in the Supporting Material**), indicating that the directionality is independent of an assay geometry and forces acting in microtubule sliding assays.

**Figure 3.**
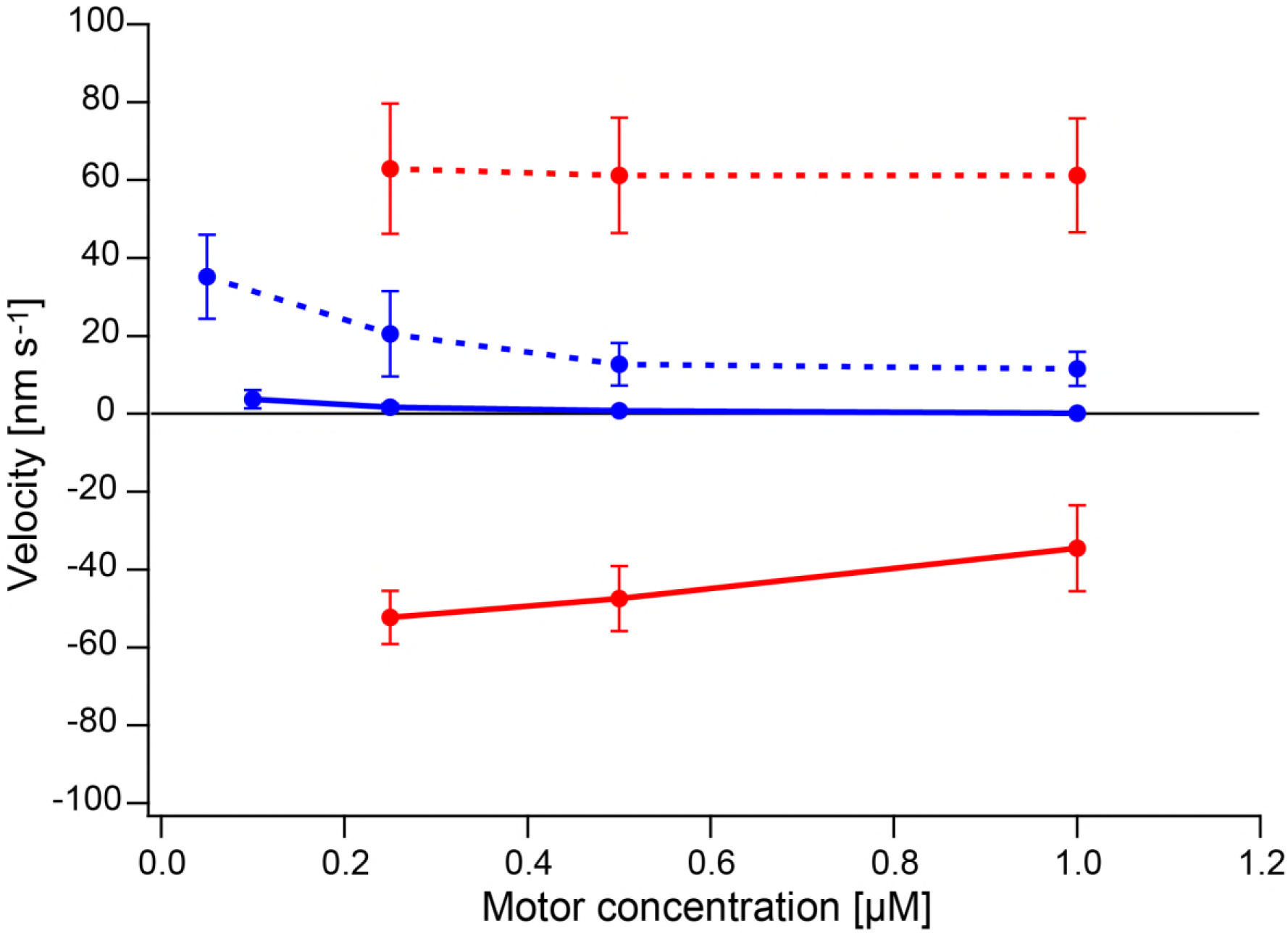
Directionalities and velocities at different motor concentrations. N-linked monomeric KlpA (red solid line) generates minus-end directionality. C-linked monomeric KlpA (red dotted line), N- and C-linked monomeric Cin8 (blue solid and dotted line) generates plus-end directionality. At concentrations below 0.25 μM for N- and C-linked KlpA, 0.1 μM for BP-Cin528 and 0.05 μM for Cin528-BP, microtubules were not observed in the sliding assay. Velocities are mean ± S.D.

**Figure 4.**
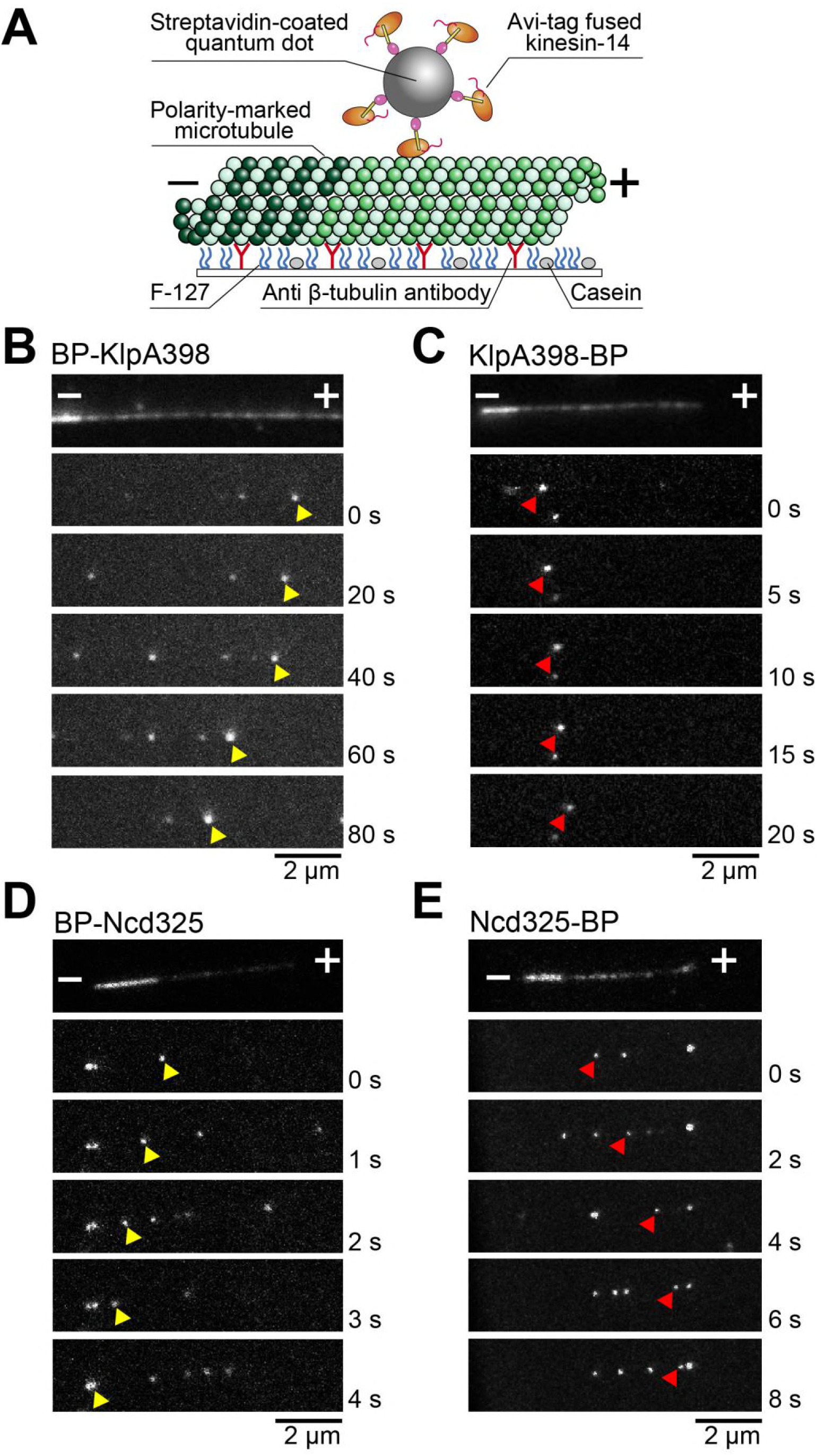
Quantum dot assay of N- or C-linked monomeric kinesin-14s. (*A*) Scheme of the quantum dot (QD) assay. Streptavidin-coated QDs, which are linked to avi-tag fused kinesin-14s via either an N- or C-terminal avi-tag and biotin-streptavidin linkage, move on polarity-marked microtubules. (*B*~*E*) Sequential images of QDs moving on polarity-marked microtubules. QD-BP-KlpA398 (*B*) and KlpA398-BP-QD (*C*) move towards the bright minus-end (yellow arrow head) and dim plus-end of the microtubule (red arrow head), respectively. Mean velocities for QD-BP-KlpA398 and KlpA398-BP-QD are -72 ± 23 nm s^−1^ (mean ± SD, n = 69) and +59 ± 17 nm s^−1^ (mean ± SD, n = 12), respectively. QD-BP-Ncd325 (*D*) and Ncd325-BP-QD (*E*) move towards the bright minus-end (yellow arrow head) and dim plus-end of the microtubule (red arrow head), respectively. Mean velocities are -30 ± 15 nm s^−1^ (mean ± SD, n = 275) and +64 ± 21 nm s^−1^ (mean ± SD, n = 141), respectively. The plus (+) and minus (-) signs refer to the plus-end and minus-end of the microtubule, respectively.

Our motility assays using minimal motor constructs show that the kinesin catalytic motor core including that of kinesin-14 defaults to plus-end directionality. The catalytic core achieves minus-end-directed motor activity by the proper function of the neck-helix cooperating with either the kinesin-14 catalytic core or neck-mimic, only when the N-terminal neck-helix is connected inline between the catalytic core and a substrate surface. Together with previous studies^2,6,7,9,11^, all C-linked kinesins, kinesin mutants and chimeras are commonly plus-end-directed even though constructs have the native neck-helix region at their N-termini (**Fig. S2**). Although the mechanism of plus-end-directed motility for the minimal kinesin motor domain remains unclear, two broad types of model have been suggested. In the first, the catalytic core itself is able to generate a small plus-end-directed conformational change on microtubules (11, 22). In the second, plus-end directionality is generated by directionally-biased binding of the catalytic core to the microtubule (23, 24). Models incorporating both processes are also possible. To carry out minus-end-directed motion, kinesin may have acquired the ability to overcome the plus-end directionality in the catalytic motor core through the introduction of the N-terminal neck-helix structure and changes in the C-terminal kinesin-1 neck-linker sequence during evolution.

## SUPPORTING MATERIAL

Two figures, two movies are available at http://www.biophsj.org/biophysj/supplemental/.

## ACKNOWLEDGEMENTS

We thank M. Sugawa for technical assistance. We are grateful to D. R. Drummond for critical reading and discussion of this manuscript. This work was supported in by JSPS KAKENHI (No.15H01629, No.15K07022 No.15H02006, No.16KT0065 to J.Y.).

## AUTHOR CONTRIBUTIONS

J.Y. designed the project. M.Y. purified the proteins and performed motility. All the authors wrote the manuscript.

## REFERENCES

1. Rice, S., A.W. Lin, D. Safer, C.L. Hart, N. Naber, B.O. Carragher, S.M. Cain, E. Pechatnikova, E.M. Wilson-Kubalek, M. Whittaker, E. Pate, R. Cooke, E.W. Taylor, R.A. Milligan, and R.D. Vale. 1999. A structural change in the kinesin motor protein that drives motility. Nature. 402: 778–84.

2. Case, R.B., D.W. Pierce, N. Hom-Booher, C.L. Hart, and R.D. Vale. 1997. The Directional Preference of Kinesin Motors Is Specified by an Element outside of the Motor Catalytic Domain. Cell. 90: 959–966.

3. Endres, N.F., C. Yoshioka, R.A. Milligan, and R.D. Vale. 2005. A lever-arm rotation drives motility of the minus-end-directed kinesin Ncd. Nature. 439: 875–878.

4. Szczęsna, E., and A. a Kasprzak. 2012. The C-terminus of kinesin-14 Ncd is a crucial component of the force generating mechanism. FEBS Lett. 586: 854–858.

5. Walker, R.A., E.D. Salmon, and S.A. Endow. 1990. The Drosophila claret segregation protein is a minus-end directed motor molecule. Nature. 347: 780–2.

6. Endow, S.A. 1998. Determinants of Kinesin Motor Polarity. Science. 281: 1200–1202.

7. Yamagishi, M., H. Shigematsu, T. Yokoyama, M. Kikkawa, M. Sugawa, M. Aoki, M. Shirouzu, J. Yajima, and R. Nitta. 2016. Structural Basis of Backwards Motion in Kinesin-1-Kinesin-14 Chimera: Implication for Kinesin-14 Motility. Structure.: 1–13.

8. Hwang, W., M.J. Lang, and M. Karplus. 2008. Force Generation in Kinesin Hinges on Cover-Neck Bundle Formation. Structure. 16: 62–71.

9. Henningsen, U., and M. Schliwa. 1997. Reversal in the direction of movement of a molecular motor. Nature. 389: 93–6.

10. Hirose, K., U. Henningsen, M. Schliwa, C. Toyoshima, T. Shimizu, M.C. Alonso, R.A. Cross, and L.A. Amos. 2000. Structural comparison of dimeric Eg5, Neurospora kinesin (Nkin) and Ncd head-Nkin neck chimera with conventional kinesin. EMBO J. 19: 5308–14.

11. Sablin, E.P., R.B. Case, S.C. Dai, C.L. Hart, A. Ruby, R.D. Vale, and R.J. Fletterick. 1998. Direction determination in the minus-end-directed kinesin motor ncd. Nature. 395: 813–6.

12. Endow, S.A. 1999. Determinants of molecular motor directionality. Nat. Cell Biol. 1: E163–7.

13. Endow, S.A., S.J. Kang, L.L. Satterwhite, M.D. Rose, V.P. Skeen, and E.D. Salmon. 1994. Yeast Kar3 is a minus-end microtubule motor protein that destabilizes microtubules preferentially at the minus ends. EMBO J. 13: 2708–13.

14. Popchock, A.R., K.-F. Tseng, P. Wang, P.A. Karplus, X. Xiang, and W. Qiu. 2017. The mitotic kinesin-14 KlpA contains a context-dependent directionality switch. Nat. Commun. 8: 13999.

15. Cull, M.G., and P.J. Schatz. 2000. Biotinylation of proteins in vivo and in vitro using small peptide tags. Methods Enzymol. 326: 430–40.

16. Yajima, J., and R. a Cross. 2005. A torque component in the kinesin-1 power stroke. Nat. Chem. Biol. 1: 338–341.

17. Heuston, E., C.E. Bronner, F.J. Kull, and S.A. Endow. 2010. A kinesin motor in a force-producing conformation. BMC Struct. Biol. 10: 19.

18. Kull, F.J., and S.A. Endow. 2013. Force generation by kinesin and myosin cytoskeletal motor proteins. J. Cell Sci. 126: 9–19.

19. Roostalu, J., C. Hentrich, P. Bieling, I.A. Telley, E. Schiebel, and T. Surrey. 2011. Directional Switching of the Kinesin Cin8 Through Motor Coupling. Science. 332: 94–99.

20. Gerson-Gurwitz, A., C. Thiede, N. Movshovich, V. Fridman, M. Podolskaya, T. Danieli, S. Lakämper, D.R. Klopfenstein, C.F. Schmidt, and L. Gheber. 2011. Directionality of individual kinesin-5 Cin8 motors is modulated by loop 8, ionic strength and microtubule geometry. EMBO J. 30: 4942–4954.

21. Britto, M., A. Goulet, S. Rizvi, O. von Loeffelholz, C.A. Moores, and R.A. Cross. 2016. Schizosaccharomyces pombe kinesin-5 switches direction using a steric blocking mechanism. Proc. Natl. Acad. Sci. U. S. A. 113: E7483–E7489.

22. Case, R.B., S. Rice, C.L. Hart, B. Ly, and R.D. Vale. 2000. Role of the kinesin neck linker and catalytic core in microtubule-based motility. Curr. Biol. 10: 157–160.

23. Grant, B.J., D.M. Gheorghe, W. Zheng, M.C. Alonso, G. Huber, M. Dlugosz, J.A. McCammon, and R.A. Cross. 2011. Electrostatically biased binding of kinesin to microtubules. PLoS Biol. 9: e1001207.

24. Okada, Y. 1999. A Processive Single-Headed Motor: Kinesin Superfamily Protein KIF1A. Science. 283: 1152–1157.

